# Synchronization between keyboard typing and neural oscillations

**DOI:** 10.1101/2020.08.25.264382

**Authors:** J. Duprez, M. Stokkermans, L. Drijvers, M.X Cohen

## Abstract

Rhythmic neural activity synchronizes with certain rhythmic behaviors, such as breathing, sniffing, saccades, and speech. The extent to which neural oscillations synchronize with higher-level and more complex behaviors is largely unknown. Here we investigated electrophysiological synchronization with keyboard typing, which is an omnipresent behavior daily engaged by an uncountably large number of people. Keyboard typing is rhythmic with frequency characteristics roughly the same as neural oscillatory dynamics associated with cognitive control, notably through midfrontal theta (4 -7 Hz) oscillations. We tested the hypothesis that synchronization occurs between typing and midfrontal theta, and breaks down when errors are committed. Thirty healthy participants typed words and sentences on a keyboard without visual feedback, while EEG was recorded. Typing rhythmicity was investigated by inter-keystroke interval analyses and by a kernel density estimation method. We used a multivariate spatial filtering technique to investigate frequency-specific synchronization between typing and neuronal oscillations. Our results demonstrate theta rhythmicity in typing (around 6.5 Hz) through the two different behavioral analyses. Synchronization between typing and neuronal oscillations occurred at frequencies ranging from 4 to 15 Hz, but to a larger extent for lower frequencies. However, peak synchronization frequency was idiosyncratic across subjects, therefore not specific to theta nor to midfrontal regions, and correlated somewhat with peak typing frequency. Errors and trials associated with stronger cognitive control were not associated with changes in synchronization at any frequency. As a whole, this study shows that brain-behavior synchronization does occur during keyboard typing but is not specific to midfrontal theta.

**Significance statement:** Every day, millions of people type on keyboards. Keyboard typing is a rhythmic behavior, with inter-keystroke-intervals of around 135 ms (~7 Hz), which is roughly the same frequency as the brain rhythm implicated in cognitive control (“theta” band, ~6 Hz). Here we investigated the hypothesis that the EEG signature of cognitive control is synchronized with keyboard typing. By recording EEG during typing in 30 healthy subjects we showed that keyboard typing indeed follows theta rhythmicity, and that synchronization between typing and neural oscillations occurs. However, synchronization was not limited to theta but occurred at frequencies ranging from 4 to 15 Hz, and in several regions. Brain-behavior synchronization during typing thus seems more nuanced and complex than we originally hypothesized.

## Introduction

In the past few decades, some sensory processes and behaviors have been shown to be rhythmic. For instance, visual processing oscillates between low and high perception stages (VanRullen, 2016), and behaviors such as speech (Aiken and Picton, 2008), although not strictly rhythmic per se, have also been associated with periodic temporal properties (Rosen, 1992). The phase of rhythmic neural activity (*i.e.* neural oscillations) has been linked to sensory and motor processes (Gross et al., 2001; Drewes and VanRullen, 2011) and some studies have reported that neural activity can become temporally aligned with rhythmic behaviors (Calderone et al., 2014; Zoefel and VanRullen, 2016; Haegens and Zion Golumbic, 2018; Kösem et al., 2018). On the other hand, neural activity can also provide temporal constraints to behaviors and several cognitive functions have been proposed to be inherently rhythmic in the sense that they are organized according to the timing of neural oscillations. Notably, this has been shown for attention (Fiebelkorn and Kastner, 2019) and cognitive control (Duprez et al., 2020).

Most of the evidence of synchronization between behavior or cognitive functioning with brain oscillatory activity have been inferred from simple experimental psychology tasks. Although the studies using such paradigms are very informative, they lack the complexity of real-life behaviors. Thus, it is important to understand whether neural oscillations and behavior synchronization extends to more naturalistic behavior. However, such naturalistic behavior should also comply with the same experimental rigor required in usual tasks, leading to difficulties in finding a good compromise between rigor and behavioral relevance.

Fortunately, technological advances also led to the emergence of newly common human behaviors. A good example of such recent and widespread behavior is keyboard typing, which is omnipresent in our modern societies, whether it is expressed on personal computers, laptops, smartphones or tablets. Studying this behavior has several advantages such as an ease in lab use (simultaneous recording of behavior and EEG with every stimulus and keypress as an event marker) and in finding performant typists. Furthermore, some evidence suggests that typing is also a rhythmic behavior with mean inter-keystroke intervals of approximately 135 ms corresponding to a 7 Hz typing frequency, which is in the theta range (Yamaguchi et al., 2013). More importantly, regarding brain activity, recent results suggest that typing has similar neural substrates to cognitive control and error monitoring. Indeed, typing errors are associated with stronger midline frontal theta power (Kalfaoğlu et al., 2018), which is commonly reported in more standard cognitive control tasks (Cavanagh and Frank, 2014; Cohen and van Gaal, 2014).

In this study, our goal was to test the hypothesis that midfrontal theta activity synchronizes with keyboard typing. We also expected that typing errors would be associated with a breakdown in this brain-behavior synchronization. To this end we used a typing task that required participants to type both real words/sentences, or pseudo-words/sentences, in which case the need for cognitive control would be greater. We first focused on showing rhythmicity in typing, and then used frequency-specific spatial filters based on a multivariate guided source separation method (generalized eigendecomposition; Cohen, 2017) to investigate synchronization between typing and neuronal oscillations.

## 1. Methods

### 2.1. Participants

Thirty healthy native Dutch speakers (16 females, 14 males, mean age: 22 years old, sd: 3.12 years) were recruited for this study through the Sona recruitment system of the Donders Institute for Brain, Cognition and Behavior, in exchange for money (20 €) or course credit. Twenty six participants were right-handed and 8 participants reported using 4 to 8 fingers instead of 10 for typing. All participants signed written informed consent and this study was approved by the Donders Institutes’ ethics committee.

### 2.2. Task

The typing task was based on the one used in Kalfaoğlu and Stafford (2014). Participants were seated in front of a 24 -inch IRT monitor and had to type the presented stimuli on a Dell qwerty-keyboard as quickly and as accurately as possible. In different blocks, we randomly presented real words, real sentences, pseudo-words, or pseudo-sentences. Participants were instructed to type what they saw without looking at the keyboard, and to press the enter key when finished. An inter-trial-interval of 2 seconds separated subsequent trials. Each block lasted 5 minutes. There were also 5 “free typing” trials during which participants responded to open questions about their activities the day before, of book/film plots, for 3 minutes with no other restrictions than typing correct Dutch sentences. These trials were not included in the analyses. No visual feedback was provided to the participants during typing. Participants were observed via a video camera and verbally instructed not to look at the keyboard when needed. The task was programmed in MATLAB using functions from psychtoolbox (Brainard, 1997) to integrate the stimuli with the hardware, and to send triggers for each keystroke to the EEG acquisition, with high temporal precision.

Participants were instructed to correct typing errors using backspace presses. They were also informed that punctuation and capital letters could be ignored. Throughout the task, the average typing speed of the participants was calculated and used as a threshold. Should they type below the threshold, the participants would be asked to type faster (except in the free typing condition) by a “Type faster” message followed by a 1s inter-trial interval. Stimuli consisted of 200 real words (average of 10.35 characters, sd = 2.5), 200 pseudo-words (average of 9.0 characters, sd = 1.8), 100 real sentences (average of 60.8 characters including spaces, sd = 13.6), and 100 pseudo-sentences (average of 59.6 characters including spaces, sd = 9.6). Pseudo-words were taken from a pseudo-word generator (Wuggy; Keuleers and Brysbaert, 2010) and sentences were taken from Drijvers et al. (2016). Pseudo-sentences were created by replacing words with pseudo-words (from the same generator) in the sentences.

### 2.3. EEG acquisition and preprocessing

EEG data were acquired at 1000 Hz using a cap with 64 active electrodes placed according to the “M10” equidistant electrode layout. Four additional electrodes were used to record horizontal and vertical eye movements. The EEG signal was amplified by BrainAmp amplifiers driven by a BrainVision Powerpack (Brain Products GmbH, Gilching, Germany). The signal was recorded using the Brain Vision software. Keystrokes were recorded as event markers in the EEG signal by sending TTL pulses. For each participant, electrode locations were recorded as Polhemus x-y-z coordinates and were further imported in eeglab (Delorme and Makeig, 2004) which was used for all further preprocessing steps. EEG data were filtered between 0.5 Hz and 40 Hz. Continuous EEG was visually inspected to identify bad channels which were removed and interpolated. 3 participants were excluded due to excessive EEG artifacts, leaving 27 participants for further analyses. After these steps, EEG was re-referenced to an average reference. Independent component analysis was performed to identify artifacts such as eye-blinks using the jader function of eeglab (Delorme and Makeig, 2004). Visual inspection of the components’ topography, activity and power spectrum was performed to identify bad components that were then removed from the signal (1.3 components on average were removed). Specific ASCII code marker numbers were used to track important events, such as stimulus display, letter keystrokes, backspace presses, and return button presses.

For all further analyses, trials were subdivided into 4 conditions: words, sentences, pseudo-words, pseudo-sentences; and into 3 trial outcomes: correct trials, corrected errors (which were defined by the occurrence of a backspace press), and other errors, which contained all other possible uncorrected errors: character omission/addition, wrong character typed, missing words in sentences, etc. Punctuation errors were not included in the analyses.

### 2.4. Behavioral analyses

#### 2.4.1. Reaction time

Reaction time (RT) was defined as the time between stimulus presentation and the first keystroke. All RTs faster than 200 ms or slower than 3 standard deviations from the mean were excluded from further analyses. RTs were then compared between conditions except free-typing (words, sentences, pseudo-words, pseudo-sentences) and outcome of the trial (correct, corrected error, other error). All statistical analyses were done using R (version 3.6.1; R Core Team, 2019). We performed linear mixed-model analyses on subject-averaged RT using the *lme* function of the *{nlme}* package (Pinheiro et al., 2020) with a 4 (conditions) + 3 (trial outcomes) design, with participant as a random intercept. We focused the analysis on subject-averaged RT because we wanted a simple model accounting for unexplained differences between participants and because modelling all trials’ RT explained less variance as compared to subject-averaged RT (based on the marginal and conditional R^2^ values). We chose not to investigate the interaction between conditions and trial outcome as further interpreting all the different contrasts would prove difficult and would not be further relevant for this study.

Graphical checks of the model’s residuals distribution were performed to ensure compliance with the model’s assumptions. For all mixed models, marginal and conditional R^2^ were calculated using the {MuMin} package (Barton, 2009) and are reported in the results. The *anova* function of the *{car}* package (Fox and Weisberg, 2019) was used on the models to extract significance of the fixed effects by the means of F-tests. In the case of significant main effects, post-hoc comparisons were performed using the *glht* function of the *{multcomp}* package that provides adjusted p-values using individual *z* tests (Hothorn et al., 2008). The significance threshold was set to p = 0.05.

#### 2.4.2. Inter-keystroke intervals

As a first method to investigate rhythmicity in typing, we focused on inter-keystroke intervals (IKI) which corresponds to the time between consecutive keystrokes. For this analysis we focused on stimuli that contained at least 4 characters. Average IKI for each condition, outcome and participant was calculated. Then, comparison between conditions and outcomes were carried out using similar linear mixed models as described for reaction time but applied on log-transformed IKI. We applied this transform because graphical check of the models’ residuals showed better compliance with the models’ assumptions when data were transformed than when left raw. We tested log, inverse, and square-root transforms and found better compliance with log based on visual inspection of the models’ residuals.

#### 2.4.3. Kernel density estimation

We quantified behavioral rhythmicity in typing by the mean of a kernel density estimation (KDE) analysis. This method is a non-parametric calculation of a probability density function. For each word, we computed the sum of the convolution of each IKI with a Gaussian kernel. The Gaussian kernel was defined as:

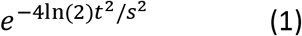

t is a time vector and s is the kernel bandwidth that was set at 1. At the IKI level, this was done by multiplying the Fourier transform of the kernel with the Fourier transform of the IKI signal, and then taking the inverse Fourier transform of the result. The IKI signal is a 0.1 Hz resolution vector ranging from 1 to 100 Hz containing only zeros except for the closest frequency of the IKI (which contains 1). For each word, the result of KDE is the sum of each IKI convolution normalized by the number of letters composing the word. The average result of KDE was calculated for each subject according to experimental conditions and outcome. Further analyses involved identifying the peak typing frequency for each condition and outcome and comparing them at the group-level. We used the same linear-mixed models as previously described. In that case, peak frequencies were log-transformed before modelling because inspection of the model’s residuals indicated better compliance with the model’s assumption in that case.

### 2.5. Spatial filtering of the data

All preprocessed EEG signal analyses were performed in Matlab (The Mathworks, USA, version 2016a) using custom written code (Cohen, 2014). In order to investigate frequency-specific synchronization between brain activity and typing behavior, we applied spatial filters constructed via generalized eigendecomposition (GED): for each frequency, a spatial filter was used to extract a single component that maximizes activity at that specific frequency. Designing the spatial filter involves computing channel covariance matrices and finding a set of channel weights that maximally differentiates the narrowband covariance from the broadband covariance (Nikulin et al., 2011; de Cheveigné and Arzounian, 2015). Calculation is done via Rayleigh’s quotient, finding a set of channel weights in vector ***w*** that maximizes λ:

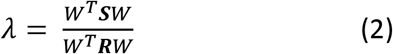

with ***S*** as the covariance matrix of the signal narrow-band filtered at the studied frequency, and ***R*** the covariance matrix from the broadband signal.

Equation (2) can be extended as a generalized eigenvalue equation:

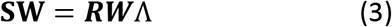

The column of **W** with the highest associated eigenvalue in Λ corresponds to the spatial filter that maximally distinguishes ***S*** from ***R***. The spatial filter can then be applied to the data as follows:

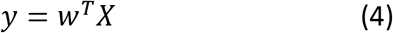

with ***w*** being the column of ***W*** corresponding to the largest associated eigenvalue, and with ***X*** the channel-by-time data matrix. The resulting component time-series is then the row vector ***y*** on which all subsequent analyses can be performed. This spatial filter method has the advantage of increasing signal-to-noise ratio, taking into account inter-individual topographical differences, and avoiding electrode selection bias (Nikulin et al., 2011; Cohen, 2017).

In this study we investigated frequencies from 3 to 15 Hz with a 0.5 Hz resolution. For each GED, narrow-band filtering of the data to create the ***S*** matrix was done using these frequencies. We limited the studied frequency to 15 Hz because we can reasonably assume that it is unlikely that one could type faster than 15 Hz frequency.

The selection of the best spatial filter ***w*** in the ***W*** result matrix was done through a combination of a statistical criterion (the size of the eigenvalues) and visual inspection of the topographical maps of the spatial filters’ activation pattern. We inspected the 15 spatial filters associated with 15 highest eigenvalues, then removed the filters that were clearly artifactual (for instance showing clear eye blink activity, or one/several electrode issues). After these steps, the component with the highest eigenvalue was chosen as the spatial filter regardless of the topographical activation pattern.

Finally, one should note that GED spatial filtering doesn’t allow us to make inferences at the cortical level and all further analyses and interpretations are limited to topographical, channel-level distributions.

### 2.6. Time-frequency decomposition

After spatial filtering of the data, time-frequency decomposition was performed to inspect time-frequency power of the components defined by GED. To this end, we used complex Morlet wavelet convolution: the Fourier transform of the data (GED component) was multiplied by a the Fourier transform of a set of complex Morlet wavelet and the taking the inverse Fourier transform of the result. Complex Morlet wavelets were defined as:

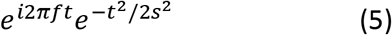

t is time, f is frequency ranging from 1 to 50 Hz (in 60 logarithmically spaced steps), and s is the width of each frequency band, defined as *n*/(2*πf*), with n increasing logarithmically from 4 to 12. We then took the squared-magnitude of the resulting signal (Z) as *real* [*Z*(*t*)^2^] + *imaginary* [*Z*(*t*)^2^] to obtain power at each frequency. Power was then baseline-corrected using a decibel (dB) transform: *dB power* = 10 × *log*10(*power*/*baseline*). Time frequency decompositions were applied either on the data epoched from −700 ms to 1500 ms around stimulus presentation. Baseline power was defined as the average power across all conditions in the period ranging from −500 to −200 ms before stimulus onset. When plotting time-frequency power results we deliberately used a different colormap than for the topographical maps of the spatial filters activation (see Figure 5C-F, and 5A-D) in order to differentiate between power and activation of the spatial filter which are completely different measures.

### 2.7. Brain-behavior synchronization measures

We tested for brain-keystroke synchronization by extracting the phase angle time series via the filter-Hilbert transform method, and computed the consistency of brain phase angles extracted from the time points of the keystrokes as follows:

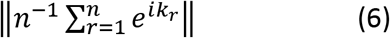

*n* is the number of keystrokes, *e*^*ik*^ is the complex representation of the phase angle *k* of keystroke *r*. Values close to 0 indicate random distribution of the phase angles, while values close to 1 indicate strong phase consistency.

At the subject level, the significance of phase consistency was calculated using the following equation:

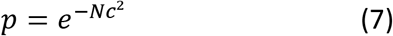

with *N* being the number of keystrokes and *c* the consistency as calculated using equation (6).

To compare phase consistency between frequency, condition and outcome, we applied a z-score normalization based on permutation analyses to the data. At each iteration, a phase angle time series vector was created using a cut-and-swap procedure that cuts the original time series at a random point and puts the second half before the first half, thus misaligning keystrokes and phases’ timing. Following this, phase consistency was calculated as described in equation (4). This procedure was carried out 500 times separately for each condition and outcome, providing a distribution of phase consistency under the null hypothesis that phase angles are randomly distributed. Z-score normalization was then applied to real consistency values by subtracting the average consistency of the null hypothesis distribution and dividing the result by its standard deviation.

To investigate frequency-specific effects of outcome and experimental condition on phase consistency, linear mixed models with a 4 (conditions) + 3 (outcomes) design, and with participant as a random effect, were performed at each frequency (using the same functions as described in section 2.4.1).

### 2.8. Data and code availability

All EEG and behavioral data will be posted on the Donders data repository upon acceptance. All the codes used for EEG analyses, statistical analyses and figures are openly available at: https://github.com/jduprez.

## 2. Results

### 3.1. Behavioral results - Is there rhythmicity in typing?

#### 3.1.1. RT

Figure 1 displays the group-average and distribution of RT showing that RT increases as a function of condition. This pattern was confirmed by the statistical analyses with a significant condition effect (F_(3, 292)_ = 217.8, p < 0.0001; conditional R^2^ = 0.91, marginal R^2^ = 0.19). Words had shorter RT (743.4 ms, sd = 132.1) than pseudo-words (756 ms, sd = 137), which had shorter RT than sentences (847.4 ms, sd = 111.5), which had shorter RT than pseudo-sentences (894.8 ms, sd = 173.6). However, this gradual increase in RT with condition was only partly supported by post-hoc analyses which showed significant differences between all conditions (p < 0.01) except for the difference between words and pseudo-words (p = 0.25). Although graphical interpretation would suggest that corrected and other errors had shorter RT than correct trials, this was not supported by the statistical analyses which showed no significant trial outcome effect (F_(2, 292)_ = 0.07, p = 0.933). As a whole, these data show an increase in RT between words/pseudo-words and sentences/pseudo-sentences.

**Figure 1:**
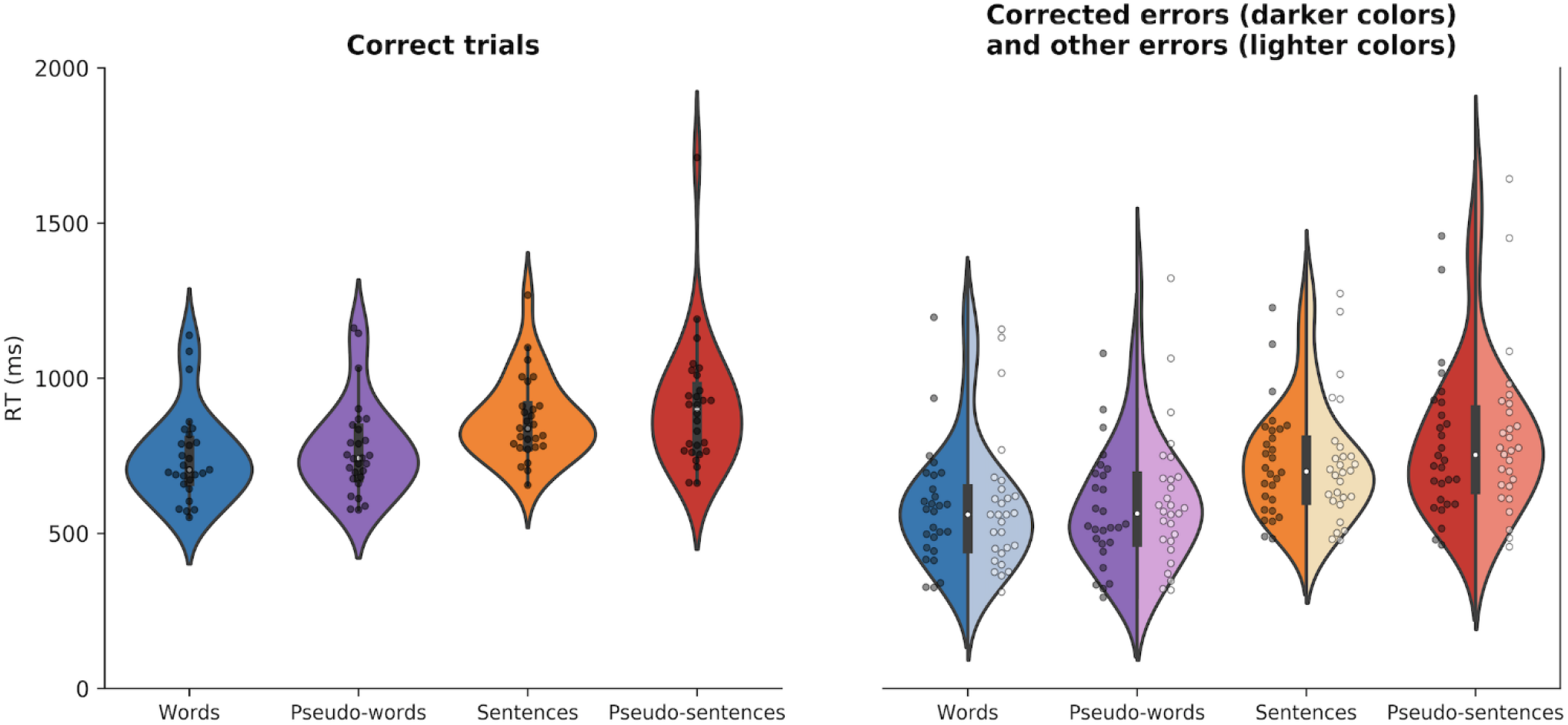
Reaction time of the first keystroke after stimulus display according to condition and trial outcome. Violin plots show the distribution of the data with an inset boxplot. Each data point corresponds to the average RT of a subject. The right part of the figure shows asymmetrical violins that present both corrected errors data (in darker colors) and other errors data (in lighter colors).

#### 3.1.2. Inter-keystroke intervals (IKI)

Overall IKI had an average of 156.2 ms (sd = 52.6), corresponding to 6.4 Hz. Figure 2 shows that IKIs changed according to condition. An increase in IKI (thus a decrease in typing frequency) can be observed between words and pseudo-words, and between sentences and pseudo-sentences. These differences were supported by the statistical analyses with a significant condition effect (F_(2, 292)_ = 93.54, p < 0.0001; conditional R^2^ = 0.84, marginal R^2^ = 0.58). Post-hoc analyses revealed significant differences between all conditions (all p < 0.001), except for the pseudo-words - pseudo-sentences comparison (p = 0.1). Words had faster IKI (162.7 ms, sd = 26) than pseudo-words (180.3 ms, sd = 29.2) and pseudo-sentences (175.8 ms, sd = 30.3), but slower IKI than sentences (152.1 ms, sd = 34.9). Figure 3 also suggests that IKI varied between trial outcomes with shorter IKI for correct trials than for corrected and other errors. We found a significant trial outcome effect (F_(2, 292)_ = 445.26, p < 0.0001) and post-hoc analyses confirmed that correct trials had shorter IKI (145.4 ms, sd = 19.5) than other errors (161 ms, sd = 24.5), which had shorter IKI than corrected errors (196.8 ms, sd = 27; all p < 0.0001). As a whole, typing frequency was faster for sentences and words compared to other conditions, and for correct trials compared to errors. It is worth noting that typing frequency was also faster for other errors than for corrected errors.

**Figure 2:**
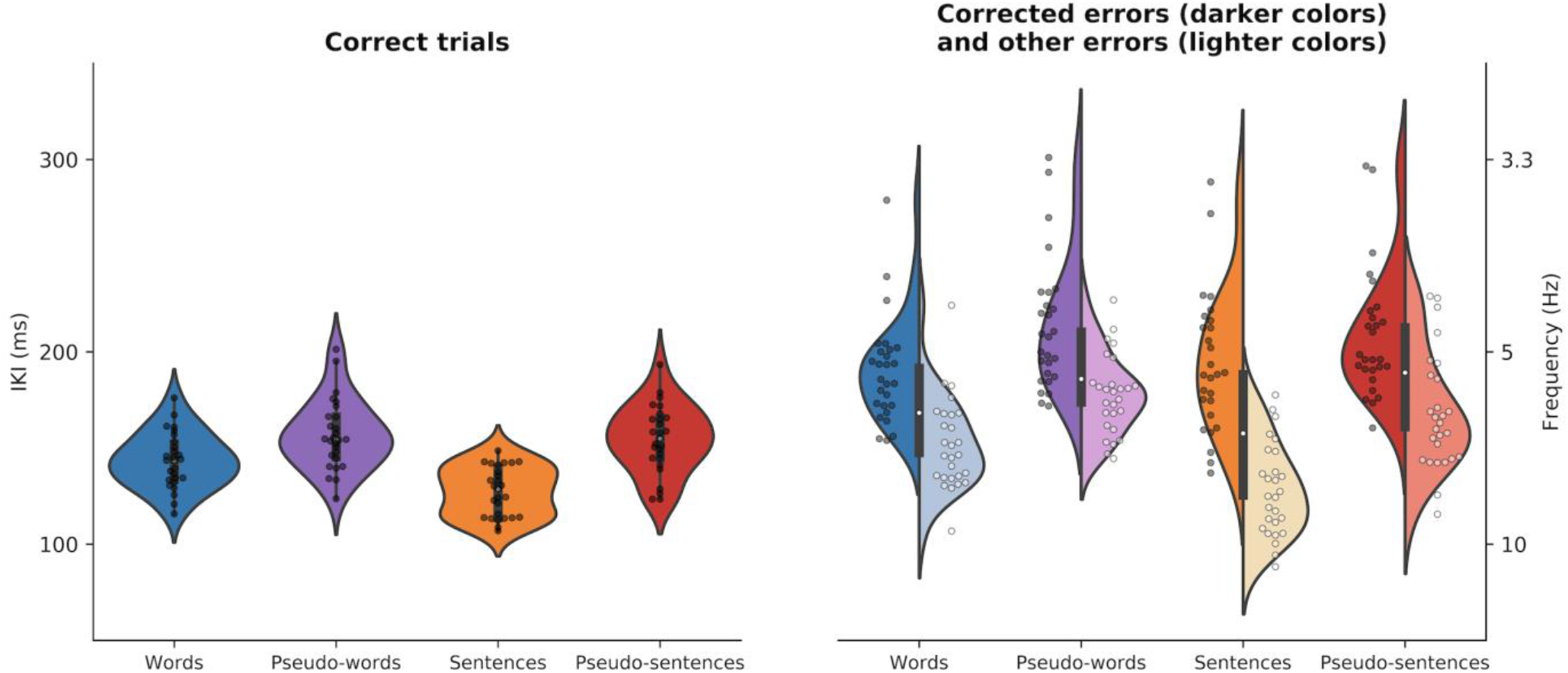
IKI according to condition and trial outcome. Violin plots show the distribution of the data with an inset boxplot. Each data point corresponds to the average IKI of a subject. The right part of the figure shows asymmetrical violins that present both corrected errors data (in darker colors) and other errors data (in lighter colors). A second y-axis has been added on the right to indicate the typing frequency corresponding to the IKI.

**Figure 3:**
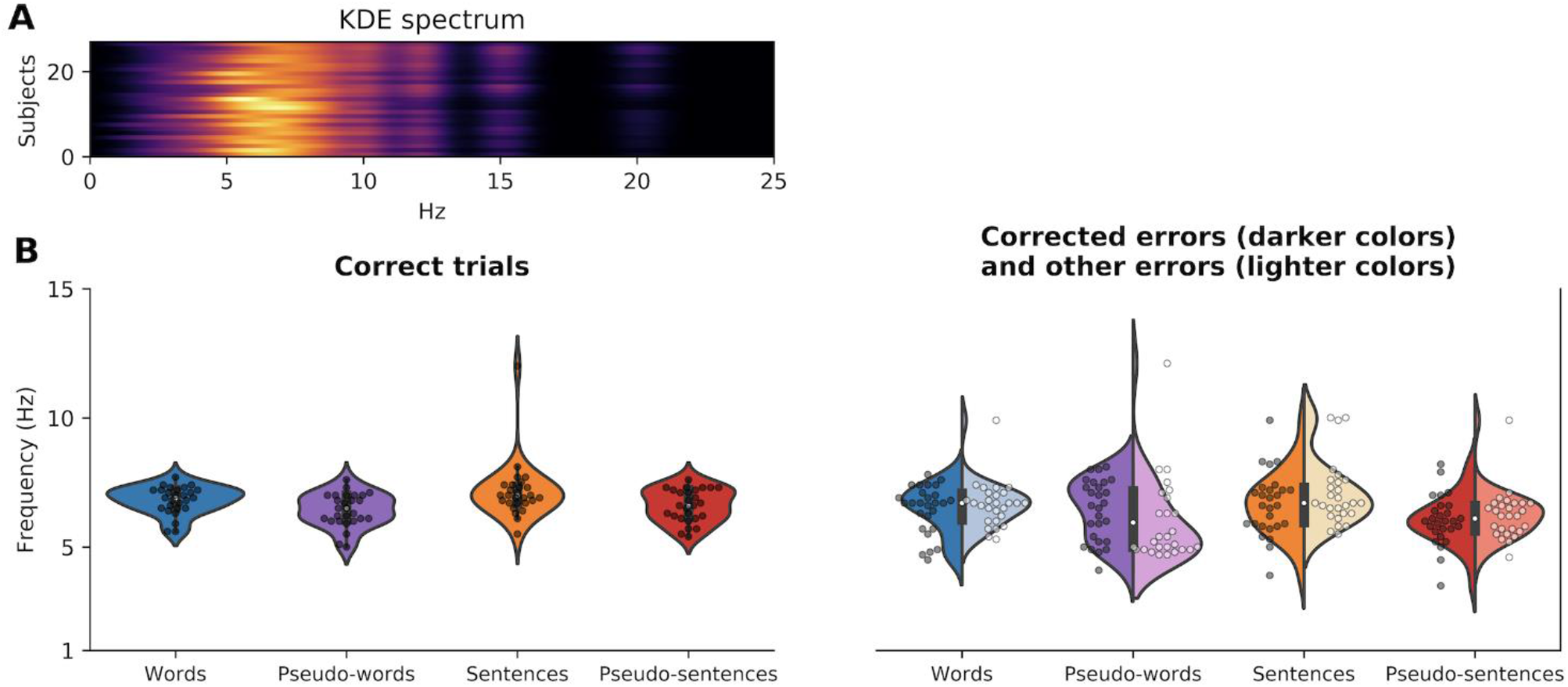
**A**: Subject specific spectrum of KDE results (averaged over all conditions for each subject - grand average of all stimuli IKI probability density function); **B**: Peak frequency extracted from the KDE result according to condition and trial outcome. Violin plots show the distribution of the data with an inset boxplot. Each data point corresponds to the peak typing frequency of a subject. The right part of the figure shows asymmetrical violins that present both corrected errors data (in darker colors) and other errors data (in lighter colors).

#### 3.1.3. Kernel density estimation (KDE) analyses

Using KDE on IKI data provided similar results as raw IKI analyses regarding typing frequency. Inspection of the KDE spectrum (Figure 3A) shows that typing mostly occurs around 6 Hz. In order to better assess typing frequency at the group level, we extracted the peak frequency for each subject according to condition and trial outcome (Figure 3B). In that case, results were less clear than when focusing on IKI. Indeed, although graphical inspection seems to show similar typing frequency regardless of condition and trial outcome, statistical analyses revealed significant differences in both effects (condition: F_(3, 286)_ = 13.85, p < 0.0001; outcome: F_(2, 286)_ = 8.33, p = 0.0003; conditional R^2^ = 0.45, marginal R^2^ = 0.09). It is worth noting that in this analysis conditional and marginal R^2^ were lower than for RT or IKI, indicating less variance explained by the model. When applying post-hoc tests, only the difference between pseudo-words and sentences was significant (p = 0.009; all other p > 0.05) with a higher frequency for sentences (6.95 Hz) than pseudo-words (6.28 Hz). Overall average peak frequency was at 6.5 Hz (sd = 1.1) which is consistent with the typing frequency obtained in the IKI results (6.4 Hz).

### 3.2. EEG - Does typing synchronize with brain oscillatory activity?

#### 3.2.1. Topographies of the spatial filters

Although we investigated frequencies ranging from 3 to 15 Hz, we only report EEG results from 4 to 15 Hz because 3 Hz activity was dominated by eye-level activity and because no participant typed at 3 Hz. Figure 4A shows the size of the largest eigenvalue according to frequencies at the group-level. The highest values were observed at 8.5 Hz which indicates that this frequency was more easily differentiated from broadband activity than other frequencies, especially from frequencies above 12 Hz. Inspection of the spatial filters created by GED revealed clear activation patterns (Figure 4B). Greater activation was centered around midfrontal electrodes from 4 to 7 Hz. A shift from midfrontal to occipital activation occurred from 8 to 11 Hz which became more occipito-parietal from 12 to 15 Hz.

**Figure 4:**
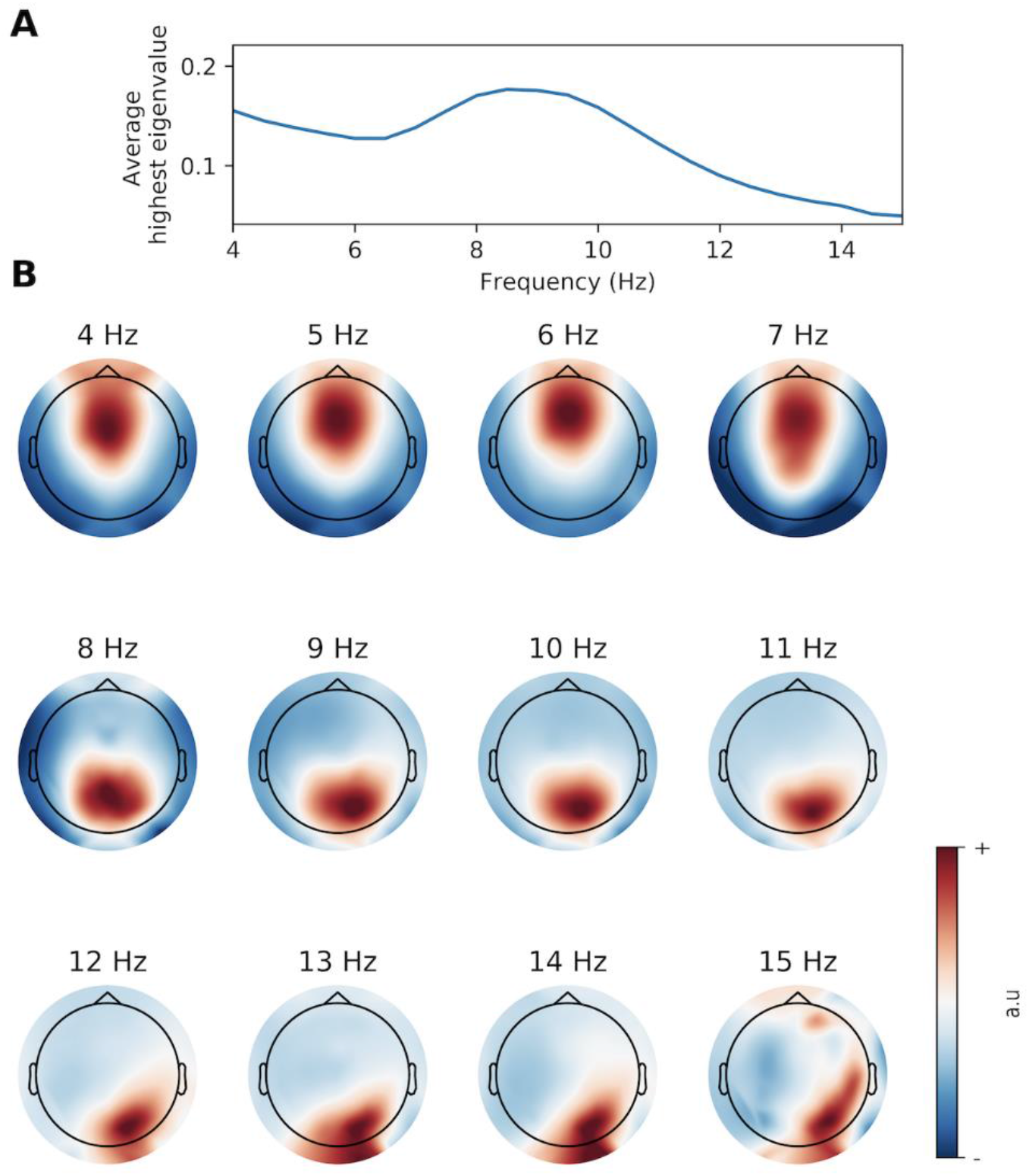
**A**: Spectrum of group-averaged highest eigenvalue. **B**: Topographical maps showing the activation pattern (in arbitrary units) of the spatial filters designed by GED for each frequency.

**Figure 5:**
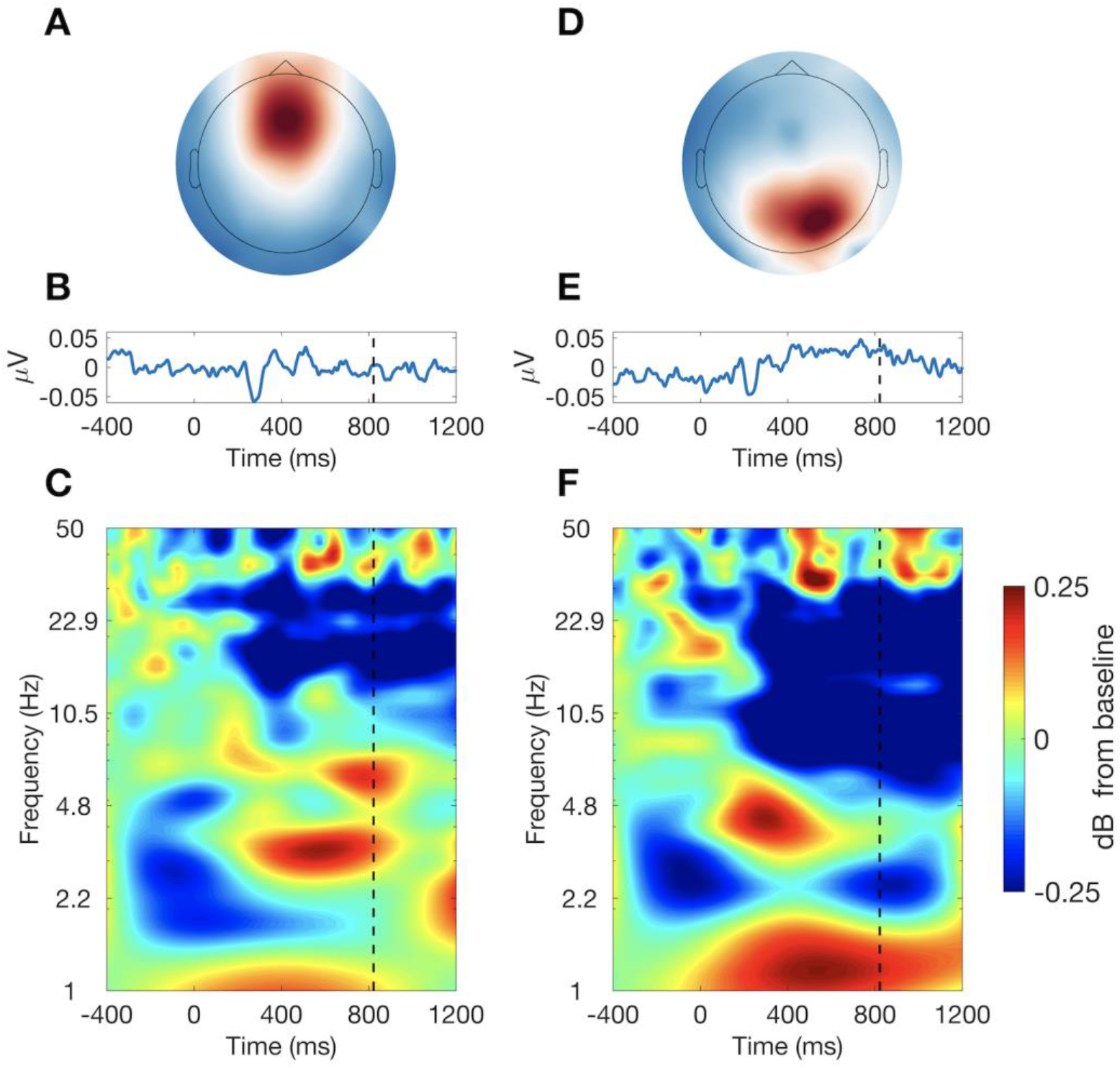
**A**: Topographical map showing the activation pattern of the 6 Hz spatial filter; **B**: Group-averaged event-related potential; **C**: Group-averaged time-frequency plot of decibel-transformed power. Stimulus onset occurred at time = 0 ms and the vertical dashed line corresponds to the group-averaged RT. **D-E-F**: same plots but for the 8.5 Hz spatial filter.

#### 3.2.2. ERP and Time-frequency analyses

In order to further evaluate the quality of the components created by GED, we inspected the ERP and time frequency decompositions of the components. Here we report only the results of the 6 Hz component, because of our hypothesis on theta synchronization with typing, and the 8.5 Hz component (Figure 5) given that it is the one associated with the highest eigenvalues. Analyzing the 6 and 8.5 Hz components revealed that the spatial filtering designed by GED resulted in physiological signals that have a clear topography, as well as time-resolved, and time-frequency characteristics. Figures 5B and 5E show the group-level ERP with clear positive and negative variations after stimulus onset.

Figures 6C and 6F reveal the time-frequency power dynamics. An increase in power can be observed between roughly 300 and 800 ms after stimulus presentation for the 6Hz component and between 200 and 600 ms for the 8.5 Hz component (thus before mean RT) in the theta band. An important decrease in power in the beta band for the 6 Hz component, and that spans the alpha and beta band for the 8.5 Hz component, was also present from roughly 300 ms to 1200 ms. Very low and high frequency changes can also be observed. The topographical map and ERP are very different from one target frequency to the other. Time-frequency power dynamics, although sharing overall similar features, also show different characteristics such as stronger alpha decrease and delta increase for the 8.5 Hz component. As a whole GED allowed to isolate frequency-specific components that each had its particular activation pattern, and oscillatory characteristics.

**Figure 6:**
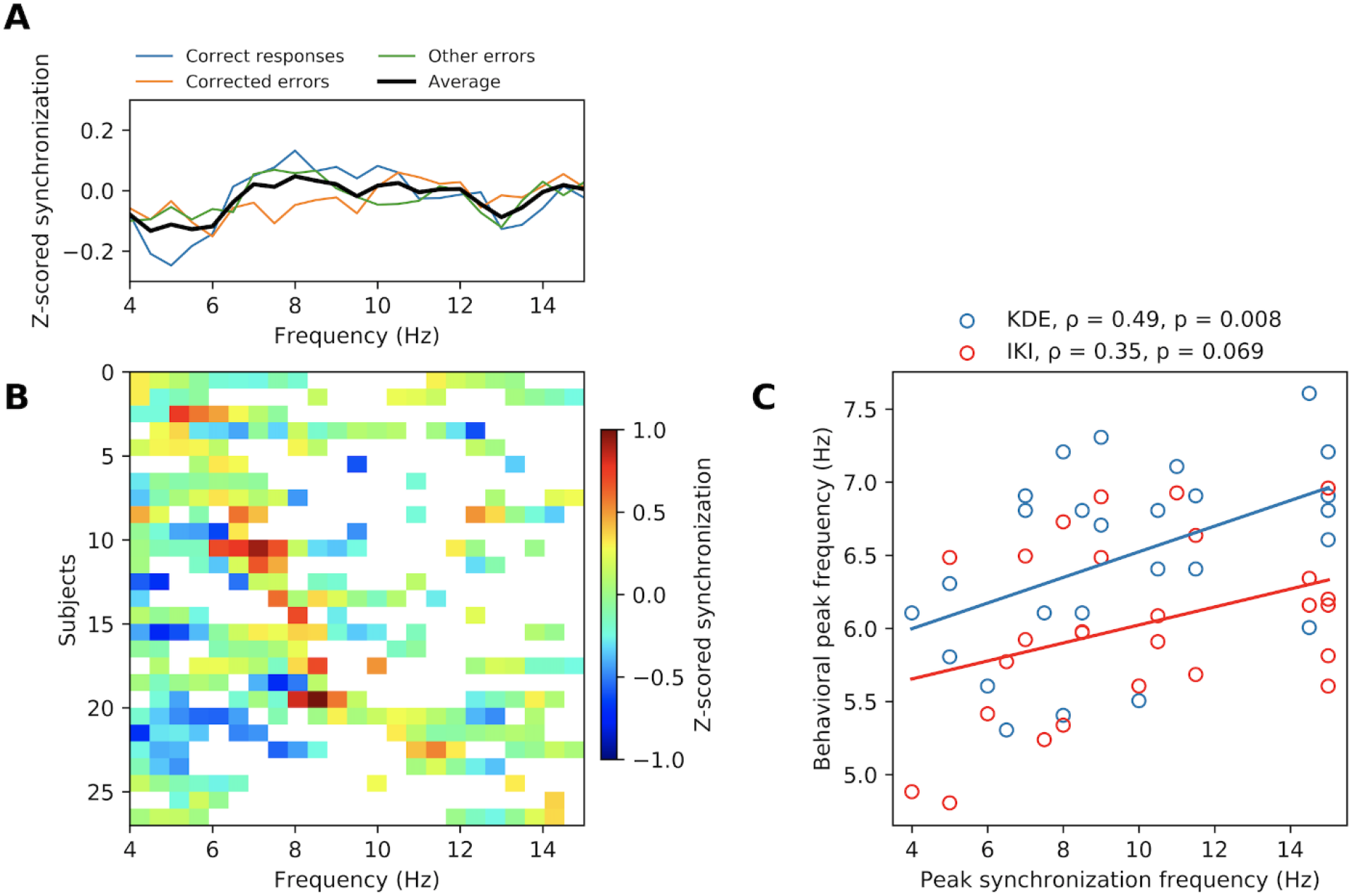
**A**: Outcome-specific and group-averaged z-scored synchronization according to frequency; **B**: Subject-specific z-scored synchronization according to frequency sorted according to peak synchronization frequency (upper subjects have a lower peak frequency). The white pixels indicate non-significant synchronization (p > 0.05) at the subject level (calculated over all trials), all other pixels have subject-level synchronization at p < 0.05; **C**: Scatterplot showing the relationship between peak synchronization frequency and behavioral peak frequency estimated by the IKI and KDE analyses.

#### 3.2.3. Frequency-specific phase clustering of all keystrokes

Our main hypothesis was that typing synchronizes with brain activity around the typing frequency. More specifically, we expected that typing in theta frequency would synchronize with midfrontal theta oscillations. For frequencies ranging from 4 to 15 Hz, phase-behavior synchronization was calculated for the corresponding GED component. Z-score normalization based on permutation analyses was then performed in order to ensure frequencies, experimental conditions, and trial outcomes are comparable.

According to Figure 6A, it seems that average synchronization between typing and brain activity peaked at 8 Hz. However, inspection of the subject-specific spectra of synchronization revealed that the peak synchronization frequency was idiosyncratic across subjects and ranged from 4 to 15 Hz (Figure 6B). Since brain-behavior synchronization could vary according to experimental conditions, we applied linear mixed models for each frequency to investigate the effect of condition and trial outcome on synchronization. This analysis didn’t reveal any significant experimental condition, or trial outcome effect at any frequency (all p > 0.05). For all frequencies conditional and marginal R^2^ were lower than 0.1 indicating only a small amount of variance was explained by the models. Furthermore, although peak synchronization frequency was highly variable across subjects, inspection of synchronization significance at the subject level revealed that most significant synchronizations occurred at frequencies lower than 9 Hz (Figure 6B). Figure 6C displays the relationship between peak synchronization frequency and behavioral peak frequency. If typing and brain activity synchronizes the most at the typing frequency, a strong correlation should appear between typing frequency and the synchronization between typing and brain activity. The correlation between peak frequency estimated through IKI analysis with the peak synchronization was not significant (ρ = 0.35, p = 0.069), whereas it was the case for correlation between the KDE-estimated frequency and peak synchronization frequency (ρ = 0.49, p = 0.008). As a whole, our results suggest that the brain-behavior synchronization mostly occurs at low frequencies during typing but is more nuanced than originally expected.

## 3. Discussion

The rhythmic nature of brain electrophysiological activity has been shown to be associated with rhythmic behaviors through various mechanisms (synchronization, entrainment, changes in behavioral performances). In this study we tested the hypothesis that keyboard typing, a real-life behavior which has been suggested to be rhythmic, synchronizes with neuronal oscillations. More specifically we expected that synchronization would preferentially occur with midfrontal theta oscillations, which have been reportedly associated with cognitive control, and recently with keyboard typing, and that this synchronization would break down during errors. Our results partly support our hypothesis by showing that typing is indeed rhythmic and that it does mostly synchronize with brain activity in low frequencies, although with no strong specificity for midfrontal theta.

### Typing is rhythmic

Keyboard typing is a highly expressed behavior in our modern societies, which makes it an ecologically valid behavior to investigate. Furthermore, studying typing is facilitated by the ease in collecting huge amounts of data in a short amount of time and without any special requirements in experimental setup and experimenter training. Typing speed has been the focus of ample research, notably regarding keyboard design and typing performance (Kinkead, 1975; Soukoreff and Mackenzie, 1995). Performances are usually assessed by investigating the number of words per minute, or through the IKI, with some studies suggesting that typing occurs in the theta frequency range (Yamaguchi et al., 2013; Kalfaoğlu and Stafford, 2014; Kalfaoğlu et al., 2018). In this study, we used two different approaches to investigate rhythmicity in typing: either by focusing on raw IKI, or by calculating their probability density function using KDE. Both approaches led to results showing that typing rhythmicity occured in the theta frequency range on average (6.4 Hz and 6.5 Hz for the IKI and KDE analyses respectively). Our results also suggest that typing reaction time and typing frequency depended to some extent on the need for cognitive control and on whether errors were committed. Indeed, pseudo-words/sentences were associated with longer RTs and lower typing frequency than real words/sentences, and the same result was observed for errors compared to correct trials (but only for typing frequency). Importantly, corrected errors showed lower typing frequency compared to other errors, which could be explained by behavioral adjustments such as post-error slowing due to error awareness and monitoring (Schroder and Moser, 2014; Ullsperger et al., 2014). However, it is important to note that these results were only found in the IKI analysis. The lack of the same effects for the KDE analysis leads to caution in interpreting condition and outcome effects on typing frequency and demands further replications.

### Unmixing frequency-specific signals

In this study we hypothesized a synchronization between typing and midfrontal theta oscillations. In order to increase the signal-to-noise ratio, to avoid bias in electrode selection, and to account for variability in subject’s topography, we chose to apply a multivariate guided source separation method, namely, generalized-eigen-decomposition (Nikulin et al., 2011; de Cheveigné and Arzounian, 2015). This method allows to create a single component that best reflects target features of the signal (in this case, a narrow-band frequency-specific signal) and has been shown to be very helpful in maximizing low-frequency features in the EEG signal (Cohen, 2017). Our results showed that this method successfully isolated different narrow-band frequency-specific components. Indeed, theta frequency components showed a midfrontal-centered activation topography, which is highly similar to topographies observed during cognitive control tasks, and were associated with comparable time-frequency dynamics such as pre-response theta power bursts (Cohen and Cavanagh, 2011; Nigbur et al., 2011; Pastötter et al., 2013; Duprez et al., 2020). Focusing on higher frequencies, topographies shifted to more posterior, occipital and occipito-parietal activations and components showed distinct time-frequency dynamics with strong alpha suppression. These topographies and power dynamics are usually associated with visual attention (Fries et al., 2008; Bauer et al., 2014; Zhang et al., 2019) and have already been reported during typing (Scaltritti et al., 2020).

### Brain-behavior synchronization during typing

Synchronization between brain activity and sensory information or behavior has been repeatedly observed in various situations. Brain oscillations can become entrained and phase-aligned to (quasi-)rhythmic stimuli with consequences on cognitive performances (see Hanslmayr et al. 2019 for an review on memory), or they can provide a more or less constrained time frame for specific cognitive processes or behavior to occur (Miller et al., 2018; Fiebelkorn and Kastner, 2019; Duprez et al., 2020). The similarities between keyboard typing and cognitive control in terms of topographical theta activity and time-frequency power (Cavanagh and Frank, 2014; Kalfaoğlu et al., 2018) led us to hypothesize the existence of synchronization between midfrontal-theta and typing behavior, and that breakdown in this synchronization would explain typing errors. However, it is important to note that our hypothesis does not presuppose whether brain oscillations would become phase-aligned with typing as a rhythmic sensory information, or whether brain oscillations would cyclically orchestrate processes resulting in rhythmic typing. Our results do not clearly support our hypothesis. Indeed, phase consistency (*i.e.* synchronization) of keystrokes did occur significantly in all subjects and at various frequencies for all subjects, suggesting that there is some broadband (non-frequency-specific) synchronization between typing and neural oscillations. Analyses showed that peak synchronization seemed to occur around 8 Hz at the group-level. Since the 8 Hz component was associated with midfrontal topography, this would support midfrontal theta synchronization with typing. However, inspection of subject-specific data showed that synchronization peak frequency was subject-specific and ranged from the 4 to 15 Hz, thus associated with components with various topographies. Therefore, although significant synchronization mostly occured at low-frequencies (and thus around midfrontal regions) at the subject-level, our results do not point to a clear specificity for the theta band and for midfrontal activity. We further reasoned that if synchronization between typing and neuronal oscillation was strong, the peak synchronization frequency and the peak typing frequency should correlate at the group-level. We only observed such correlation when focusing on typing frequency determined by the KDE method.

Moreover, no significant changes in phase consistency were observed between the different experimental conditions, and between the different trial outcomes, at any frequency. This suggests that the need for cognitive control did not modulate synchronization and that erroneous trials (whether they were corrected or not) were not necessarily associated with a decrease in synchronization.

To summarize, although our results show synchronization between typing and neuronal oscillations, it does not seem to be limited to the theta band or to midfrontal regions, and typing behavior does not seem to depend on the strength of this synchronization.

### Limitations and perspectives

This study is the first, to our knowledge, to investigate synchronization between brain activity and typing. As a result, interpretations should remain cautious and take into account some limitations.

Our results show that there was a clear variability in typing frequency across participants with some subjects showing 4.5 Hz typing frequency whereas others went up to 7.5 Hz. Such differences could participate in enhancing variability when investigating synchronization. Focusing on a group of subjects sharing more similar typing frequencies could help in diminishing inter-individual typing differences. However, it is important to note that our results show that even for similar typing frequencies, the peak synchronization frequency varied from 4 to 15 Hz. This suggests that typing frequency variability probably does not entirely account for synchronization differences.

With this study we aimed at testing a specific hypothesis on midfrontal theta synchronization with typing. Thus, we chose to use a spatial filtering technique (GED) specifically designed to enhance narrow-band frequency-specific features of the signal. Although GED successfully isolated target components, it also potentially masked broader topographical networks, which do not necessarily operate in narrow-band frequencies but rather by the means of cross-frequency dynamics. An interesting perspective of this work could be to investigate such cross-frequency networks by using subject-specific peak synchronization frequency as the low frequency timing higher frequency characteristics.

Another aspect to bear in mind is that this study was limited to the investigation of cortical brain-behavior synchronization. The synchronization we expected might be more prominent between typing and activity in the basal ganglia or the cerebellum, which are hardly accessible through EEG. For instance, in the case of cognitive control, theta oscillations in the subthalamic nucleus have been associated with conflict resolution and inhibition and subtends functional connectivity with midfrontal cortical regions (Zavala et al., 2015), making it an interesting candidate structure for synchronization with typing. Another potential structure for synchronization might be the cerebellum, which is involved in the sub-second timing of voluntary movements (Bareš et al., 2019), thus in the frequency range of keyboard typing. Finally, given the complex interplay between cortical regions, the basal ganglia and the cerebellum for goal-directed behavior, it is highly unlikely that synchronization would be exclusive to one structure. Comparing the strength of synchronization between these brain regions and typing would therefore provide interesting insights on the role of brain-behavior synchronization during keyboard typing.

## Acknowledgments

JD was funded by the Rennes Clinical Neuroscience Institute (INCR: www.incr.fr). MXC is funded by ERC-StG 638589. LD is funded by ERC-COG 773079.

## Notes

### Competing Interest Statement

The authors have declared no competing interest.

## 4. References

Bareš M, Apps R, Avanzino L, Breska A, D’Angelo E, Filip P, Gerwig M, Ivry RB, Lawrenson CL, Louis ED, Lusk NA, Manto M, Meck WH, Mitoma H, Petter EA (2019) Consensus paper: Decoding the Contributions of the Cerebellum as a Time Machine. From Neurons to Clinical Applications. Cerebellum Lond Engl 18:266–286.

Barton K (2009) MuMIn: multi-model inference. Httpr-Forge R-Proj Orgprojectsmumin.

Bauer M, Stenner M-P, Friston KJ, Dolan RJ (2014) Attentional Modulation of Alpha/Beta and Gamma Oscillations Reflect Functionally Distinct Processes. J Neurosci 34:16117–16125.

Brainard DH (1997) The psychophysics toolbox. Spat Vis 10:433–436.

Calderone DJ, Lakatos P, Butler PD, Castellanos FX (2014) Entrainment of neural oscillations as a modifiable substrate of attention. Trends Cogn Sci 18:300–309.

Cavanagh JF, Frank MJ (2014) Frontal theta as a mechanism for cognitive control. Trends Cogn Sci 18:414–421.

Cohen MX (2014) Analyzing neural time series data: theory and practice. MIT press.

Cohen MX (2017) Multivariate cross-frequency coupling via generalized eigendecomposition. ELife 6.

Cohen MX, Cavanagh JF (2011) Single-trial regression elucidates the role of prefrontal theta oscillations in response conflict. Front Psychol 2:30.

Cohen MX, van Gaal S (2014) Subthreshold muscle twitches dissociate oscillatory neural signatures of conflicts from errors. NeuroImage 86:503–513.

de Cheveigné A, Arzounian D (2015) Scanning for oscillations. J Neural Eng 12:066020.

Delorme A, Makeig S (2004) EEGLAB: an open source toolbox for analysis of single-trial EEG dynamics including independent component analysis. J Neurosci Methods 134:9–21.

Drewes J, VanRullen R (2011) This Is the Rhythm of Your Eyes: The Phase of Ongoing Electroencephalogram Oscillations Modulates Saccadic Reaction Time. J Neurosci 31:4698–4708.

Drijvers L, Mulder K, Ernestus M (2016) Alpha and gamma band oscillations index differential processing of acoustically reduced and full forms. Brain Lang 153-154:27–37.

Duprez J, Gulbinaite R, Cohen MX (2020) Midfrontal theta phase coordinates behaviorally relevant brain computations during cognitive control. NeuroImage 207:116340.

Fiebelkorn IC, Kastner S (2019) A Rhythmic Theory of Attention. Trends Cogn Sci 23:87–101.

Fox J, Weisberg S (2019) An R Companion to Applied Regression, Third. Thousand Oaks CA: Sage. Available at: https://socialsciences.mcmaster.ca/jfox/Books/Companion/.

Fries P, Womelsdorf T, Oostenveld R, Desimone R (2008) The Effects of Visual Stimulation and Selective Visual Attention on Rhythmic Neuronal Synchronization in Macaque Area V4. J Neurosci 28:4823–4835.

Gross J, Kujala J, Hämäläinen M, Timmermann L, Schnitzler A, Salmelin R (2001) Dynamic imaging of coherent sources: Studying neural interactions in the human brain. Proc Natl Acad Sci U S A 98:694–699.

Haegens S, Zion Golumbic E (2018) Rhythmic facilitation of sensory processing: A critical review. Neurosci Biobehav Rev 86:150–165.

Hanslmayr S, Axmacher N, Inman CS (2019) Modulating Human Memory via Entrainment of Brain Oscillations. Trends Neurosci 42:485–499.

Hothorn T, Bretz F, Westfall P (2008) Simultaneous Inference in General Parametric Models. Biom J 50:346–363.

Kalfaoğlu Ç, Stafford T (2014) Performance breakdown effects dissociate from error detection effects in typing. Q J Exp Psychol 2006 67:508–524.

Kalfaoğlu Ç, Stafford T, Milne E (2018) Frontal theta band oscillations predict error correction and posterror slowing in typing. J Exp Psychol Hum Percept Perform 44:69–88.

Keuleers E, Brysbaert M (2010) Wuggy: a multilingual pseudoword generator. Behav Res Methods 42:627–633.

Kinkead R (1975) Typing speed, keying rates, and optimal keyboard layouts. In: Proceedings of the Human Factors Society Annual Meeting, pp 159–161. SAGE Publications Sage CA: Los Angeles, CA.

Kösem A, Bosker HR, Takashima A, Meyer A, Jensen O, Hagoort P (2018) Neural Entrainment Determines the Words We Hear. Curr Biol CB 28:2867–2875.e3.

Miller EK, Lundqvist M, Bastos AM (2018) Working Memory 2.0. Neuron 100:463–475.

Nigbur R, Ivanova G, Stürmer B (2011) Theta power as a marker for cognitive interference. Clin Neurophysiol Off J Int Fed Clin Neurophysiol 122:2185–2194.

Nikulin VV, Nolte G, Curio G (2011) A novel method for reliable and fast extraction of neuronal EEG/MEG oscillations on the basis of spatio-spectral decomposition. NeuroImage 55:1528–1535.

Pastötter B, Dreisbach G, Bäuml K-HT (2013) Dynamic adjustments of cognitive control: oscillatory correlates of the conflict adaptation effect. J Cogn Neurosci 25:2167–2178.

Pinheiro J, Bates D, DebRoy S, Sarkar D, R Core Team (2020) nlme: Linear and Nonlinear Mixed Effects Models. Available at: https://CRAN.R-project.org/package=nlme.

Rosen S (1992) Temporal Information in Speech: Acoustic, Auditory and Linguistic Aspects. Philos Trans Biol Sci 336:367–373.

Scaltritti M, Suitner C, Peressotti F (2020) Language and motor processing in reading and typing: Insights from beta-frequency band power modulations. Brain Lang 204:104758.

Schroder HS, Moser JS (2014) Improving the study of error monitoring with consideration of behavioral performance measures. Front Hum Neurosci 8 Available at: https://www.frontiersin.org/articles/10.3389/fnhum.2014.00178/full [Accessed June 30, 2020].

Soukoreff RW, Mackenzie IS (1995) Theoretical upper and lower bounds on typing speed using a stylus and a soft keyboard. Behav Inf Technol 14:370–379.

Ullsperger M, Fischer AG, Nigbur R, Endrass T (2014) Neural mechanisms and temporal dynamics of performance monitoring. Trends Cogn Sci 18:259–267.

VanRullen R (2016) Perceptual Cycles. Trends Cogn Sci 20:723–735.

Yamaguchi M, Crump MJC, Logan GD (2013) Speed-accuracy trade-off in skilled typewriting: decomposing the contributions of hierarchical control loops. J Exp Psychol Hum Percept Perform 39:678–699.

Zavala B, Zaghloul K, Brown P (2015) The subthalamic nucleus, oscillations, and conflict. Mov Disord 30:328–338.

Zhang Y, Zhang Y, Cai P, Luo H, Fang F (2019) The causal role of α-oscillations in feature binding. Proc Natl Acad Sci 116:17023–17028.

Zoefel B, VanRullen R (2016) EEG oscillations entrain their phase to high-level features of speech sound. NeuroImage 124:16–23.

